# Growth rate and adaptive capacity, not just thermal tolerance, are critical for long-term coral persistence under climate change

**DOI:** 10.1101/2025.09.24.678233

**Authors:** Noam S. Vogt-Vincent, Mariana Rocha de Souza, Kathryn Feloy, Robert J. Toonen, Lisa C. McManus

## Abstract

It is widely assumed that corals with a narrower thermal tolerance and greater bleaching susceptibility are at most risk under anthropogenic climate change. Using a simple eco-evolutionary model, we investigate coral reef futures under climate change through a factorial approach, across millions of simulations. We find that, while corals with low thermal tolerance are indeed at greatest risk of population collapse in the short term, coral population persistence beyond 2050 is instead primarily determined by the maximum population growth rate and additive genetic variance. Anthropogenic climate change exceeds the thermal tolerance of all corals, so adaptive capacity and the ability to rapidly recover from disturbance are critical for long-term persistence. As a result, it is plausible that branching corals could fare better than slower growing, stress-tolerant counterparts in the long term, but only if they can weather the greater acute threats over the coming decades. Since the response timescale of corals to climate change varies considerably across taxa, and due to the particular importance of genetic variance in facilitating evolutionary adaptation, sustainable restoration approaches should aim to conserve a broad range of coral species and genotypes, rather than aiming to optimise one particular trait.

## Introduction

Coral reefs are among the most biologically diverse and productive ecosystems on Earth, providing valuable services such as tourism, coastal protection, natural products, and nutrition. Despite their ecological importance, coral reefs are in decline (1). The increase in temperature caused by anthropogenic climate change leads to coral bleaching, in which reef-building corals lose the algal symbionts that provide up to 90% of a corals’ energy (2,3). Bleaching results in decreased coral growth and reproductive output, as well as increased susceptibility to disease and mortality (4–6). Over the past 100 years, ocean temperatures in many tropical regions have increased by almost 1°C and are projected to rise by 1-2 °C per century under the best-case emission scenarios (7). These future conditions are predicted to have bleak outcomes for coral reefs, including potential global coral cover declines of up to 99% if temperature rises 2°C by the end of the century (10, but see 11,12).

Several factors influence the persistence of coral populations in the face of increasingly frequent and severe marine heatwaves (11). These include additive genetic variance, which supports adaptive potential (9,12); population growth rate (13,14); thermal tolerance (1,15); fecundity (16); larval immigration and connectivity (17,18); and thermal optima, which define the temperature ranges within which corals can successfully function (19,20). Coral persistence is commonly assessed through changes in reef cover, a metric shaped by key demographic and physiological processes such as survival, reproduction, and growth.

Reef recovery following a bleaching event depends heavily on the frequency, growth, and reproduction of resilient genotypes, as well as the connectivity among populations. Successful reproduction produces resilient larvae that can repopulate affected reefs, increasing the proportion of heat-tolerant individuals. These larvae can also disperse to other reefs, boosting resilience in degraded areas, and enabling the exchange of resilient genotypes between reefs. Strong evidence highlights the critical role of biological and ecological connectivity in replenishing populations and supporting ecosystem recovery after disturbances (21–23).

Despite significant research on this topic (24–27) the relative importance of factors such as coral growth rates, genetic variance, thermal tolerance, and their interactions in determining coral persistence under global change remains poorly understood. Identifying which of these factors most strongly influence persistence is critical for guiding targeted and effective management and conservation strategies. To address this gap, we applied an eco-evolutionary model based on (10,28) to explore how these factors shape coral persistence under projected future climate scenarios. While many studies have used eco-evolutionary models to project outcomes for specific species (29–31) or sites (21,32,33), our approach is not constrained to species types or locations. Instead, it identifies the most influential factors driving coral persistence, offering insights applicable across diverse reef systems. The model integrates empirical biological and environmental data to assess coral sensitivity to these key parameters. While we use Hawaiian reefs as a representative case study, the framework and findings are broadly relevant and can be extended to coral reef ecosystems globally.

Our goal is to identify the biological and ecological parameters that most strongly determine coral persistence under projected climate change. To do so, we comprehensively explore the parameter space that characterizes modern coral species. Additionally, we examine the role of larval immigration—both natural and restoration-driven—by testing how varying immigration rates influence long-term coral cover, offering insights into the potential efficacy of restoration interventions.

## Methods

In this study, we use a simple, single-population eco-evolutionary model. We model coral cover assuming that coral colony size distribution is fixed and log-normal, and that the colony linear growth rate has a maximum at a certain thermal optimum and falls as the temperature deviates from the thermal optimum, with thermal tolerance breadth *w*. Above the thermal optimum, the mortality rate rises quadratically due to thermal stress and bleaching. Colony-specific thermal optima are distributed around a population mean trait value *z* with a constant additive genetic variance *V*. In addition to growth through linear extension (i.e. budding), coral cover can also increase through larval recruitment, either from local larvae or immigrants. The population mean trait value *z* can evolve through time based on selection (depending on the sensitivity of the growth rate to the trait value *z*) or the trait values of immigrant larvae.

In all our experiments, coral cover is first allowed to equilibrate under control temperature forcing, before imposing projected 21^st^ century warming from a dynamically downscaled ocean model for the Main Hawaiian Islands (34). Due to the simplicity of the model, we can comprehensively investigate the sensitivity of coral cover dynamics to the imposed warming across parameter space. The modelling and experimental frameworks are described in further detail below.

## Modelling framework

### Ecological dynamics

For simplicity, we consider a single coral population that can receive external larval input, and do not consider inter-species interactions. Following (28,35), we assume that the fractional coral cover *N* based on local population dynamics, genetic load, and larval recruitment,

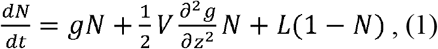

where *g*(y^-1^) is the intrinsic population growth rate, *V* (K^2^) is the additive genetic variance, *z* is the population mean thermal optimum (K), *f* is the population growth rate due to self-recruitment (y^-1^), and *L* is the larval supply (y^-1^). We decompose the growth rate *g* into a (logistic) positive growth rate *r(1 - N)* and mortality rate *m*, i.e.

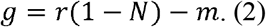

Similarly to (28), we assume that the positive growth rate falls smoothly as the temperature *T* (K) deviates from the population mean thermal optimum *z*, such that

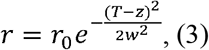

Where *r*_*0*_ is the optimal population growth rate (y^-1^) and *w* is the coral thermal tolerance (K). As in (10), we further assume that the mortality rate increases quadratically above the coral thermal optimum:

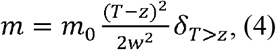

where *m*_*0*_ is a mortality rate scaling factor (y^-1^) and *δ*_*T*>*z*_ *= 1* if *T > z* and 0 otherwise.

The larval supply *L* is computed at the start of an annual spawning event as *L = fN+ I*, where *f* is an effective fecundity (y^-1^) representing self-recruitment, and *I* represents immigration (y^-1^).

### Evolutionary dynamics

Following (28), we assume that the population mean thermal optimum *z* can evolve based on selection and the trait values of immigrant larvae. After each spawning event, we compute the new value of *z* based on the number of settling immigrant larvae, i.e.

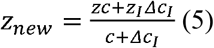

where *z*_*I*_ is the mean thermal optimum among immigrant larvae, and *Δc*_*I*_ is the number of settling immigrant larvae. At all time-steps, the effect of selection is modelled as follows:

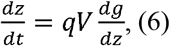

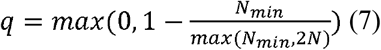

where *q* is a term that reduces the rate of selection at very small population sizes (∼*10*^*-*3^). For simplicity, we do not model interactive external larval sources and assume that the offset *z*_*I*_ *-z* is fixed, i.e. all reefs are evolving at the same rate. This assumption is justified given the restricted range of warming rates within shallow-water reef systems (34).

### Parameterization

To investigate the sensitivity of coral population dynamics under climate change to coral traits, it is important to establish a plausible parameter space. Here, we do not attempt to realistically model the exact distribution of coral traits, but rather define a plausible (order-of-magnitude) parameter envelope. We do not consider covariance between parameters, although acknowledge that these parameters may not be independent in reality (36).

### Optimal population growth rate

The optimal population growth rate *r*_0_ (y^-1^) is a function of the maximum colony linear extension rate *ε* m y^-1^ and the population colony size (planar area) structure, which we assume is log-normal with log-transformed mean and standard deviation *µ* and *σ* (37). Following (10), if colonies have a circular planar area and do not overlap,

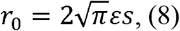

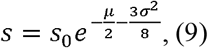

where *s* is a colony size distribution factor, *µ* and *σ* describe the log-normal planar area distribution of coral colonies, and *s*_0_ is a dummy variable with value 1 m^-1^. We do not permit this size structure to evolve through time, assuming that demographic processes keep the size structure approximately fixed. Coral colony maximum linear extension rate is typically 0.5 - 10 cm yr^-1^ (36) and the size distribution factor typically varies between 0.5 - 4 m^-1^ (37). These values are broadly log-normally distributed and, taking the above ranges to represent 1*σ* and assuming they are independent, the resulting optimal population growth range is c. 0.02 - 0.6 y^-1^.

### Mortality rate scaling factor

As in (10), we simplistically assume that mortality rate scales quadratically with temperature as the temperature exceeds the thermal optimum. In this case, the proportion of corals *N/N*_0_ that survive after one time-step *Δt* (y) is given by

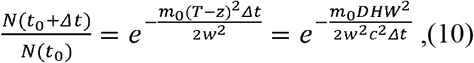

defining one degree-heating-week (DHW) as *DHW = (T - z)cΔt, c = 52* weeks y^-1^. Laboratory heat stress experiments indeed show that coral survivorship plausibly follows a similarly shaped dependence on degree-heating-weeks (11,38). Defining *DHW*_*S0*_ as the thermal stress required to cause 50% mortality and using the model time-step 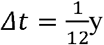, we find that

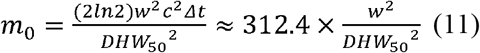

We therefore parameterise the mortality rate scaling factor based on the thermal tolerance breadth *w* and *DHW*_*S0*_, which we explore in the range 4 - 24 °C-weeks based on values reported in the literature, acknowledging that there are few known corals with thermal tolerance at the upper end of this range (11,38).

### Thermal tolerance breadth

This parameter, *w* (K), defines how the coral growth rate declines as the temperature deviates from the thermal optimum, and plausibly varies between 2 - 8 °C (19,39,40).

### Additive genetic variance

The additive genetic variance, *V* (C^2^), is the product of the narrow-sense heritability and phenotypic variance of the coral thermal optimum *z*. Quantitative trait values associated with coral thermal tolerance tend to have a heritability in the range 0.25-0.5 (9,41). Phenotypic variance in thermal tolerance thresholds within reefs have been measured at between 0.05 - 1.0 °C^2^ (42,43). This results in a plausible additive genetic variance range of 0.01 - 0.5 °C^2^.

### Effective fecundity

The effective fecundity, *f* (y^-1^), represents the potential for coral cover to increase based on self-recruitment. As in (10), we set

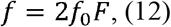

where *f*_0_ is the number of eggs produced per year (or spawning event) per polyp (y^-1^) and *F* is the likelihood that an egg transitions into a sexually mature coral within the same reef. This assumes that coral colonies are (i) hemispheric and (ii) have the size of a single coral polyp at the point of sexual maturity. We further set

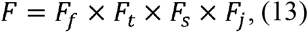

where *F*_*f*_ is the proportion of eggs that are fertilised and develop into larvae, *F*_*t*_ is the proportion of larvae that remain at the parent reef, *F*_*s*_ is the proportion of larvae remaining at the parent reef that successfully undergo recruitment, and *F*_*j*_ is the proportion of recruits that survive to sexual maturity. To obtain an order-of-magnitude estimate, we set *F*_*f*_ = [0.01,1] (44), *F*_*t*_ = [0.01,1] based on coral larval planktonic larval durations and retention rates in different coral reef systems (45,46), *F*_*s*_ = [0.01,1] (47,48), and *F*_*j*_ *=* [0.01,0.2] (48,49). The polyp-specific fecundity *f*_0_ varies enormously across the range [1,1000] (50). Assuming these variables are independent and log-normally distributed, this results in a plausible (1*σ*) range of around = [0.0001, 0.4] y^-1^, although we consider values at the lower end to be unlikely.

### Immigration

The immigration rate *I* has the same units as *f*, although is not multiplied by *N*. In this study, we keep *I* fixed in time (thereby representing a steady larval supply) and test values in the range *I =* [0.0001,0.1], comparable to, but slightly lower than, the range of plausible values for self-recruitment.

### Immigrant thermal optimum

Based on the diversity of thermal environments in reef systems such as those in Hawai□i (51) and the potential for strong intra and inter-reef connectivity, particularly for broadcast spawning corals (52,53), it is plausible that immigrant larvae may have significantly different thermal optima to the local coral population. Based on the range of mean temperatures found in Kāne□ohe Bay (51), we explore immigrant thermal optima *z*_*I*_ that are in the range [-0.5, 0.5] °C relative to the local thermal optimum *z*.

### Experimental design

To investigate the response of hypothetical coral populations to future anthropogenic climate change, we impose monthly temperatures (2002 to 2100) averaged across the 1 - 30 m depth range from a 4 km resolution ocean model for the Main Hawaiian Islands (34). These oceanographic simulations reasonably reproduce observed present-day (2000-2020) sea-surface temperature and variability (34) and projections are dynamically downscaled from the CESM2 contribution to CMIP6 (54), which is characterised by a moderate transient climate response and high equilibrium climate sensitivity among CMIP6 models (55). We use projections based on three emissions scenarios: SSP1-2.6, SSP2-4.5, and SSP3-7.0. For coastal environments in the Main Hawaiian Islands, these scenarios are associated with a modelled 21^st^ century warming trend of around 1.7 °C, 2.2 °C, and 3.1 °C respectively (figure S1). While there is relatively little spatial variability in the projected long-term warming trend within the Main Hawaiian Islands, temperature variability differs across the island chain (34). We therefore use temperature time-series from seven representative sites across the islands of Kaua□i, O□ahu, L□na□i, and Hawai□i Island that capture the range of interannual, seasonal, and sub-seasonal temperature variability within the Main Hawaiian Islands (figures S1-2).

In each eco-evolutionary simulation, we initialise coral cover at the carrying capacity and impose repeating temperature forcing based on 2002-2011 monthly averages to allow model equilibration to these initial conditions. After verifying that equilibrium has been reached, we impose monthly temperatures separately for each of the seven sites and three emissions scenarios.

We investigate the sensitivity of coral cover trajectories to coral biological and ecological parameters through a factorial sensitivity analysis. We test seven values across the stated range for each of the seven variables in Table 1, for a total of 5.8 million individual simulations for each of the three emissions scenarios. To compute 21^st^ century ‘coral cover decline’, we calculate decadal coral cover averages to remove the effects of short-term variability, and then calculate the change in coral cover between the first ten years of the simulation and the decade with the lowest coral cover. Since different parameter combinations may be associated with different equilibrium initial conditions, we base our analyses on the relative, rather than absolute, change in coral cover. Changes in coral cover in this manuscript always refer to relative changes unless otherwise noted. We quantify the sensitivity of change in coral cover to model parameters using random forest regression and permutation importance (56). The permutation importance of a certain parameter gives the reduction in the R^2^ score from the random forest regressor when predictor values from that parameter are randomly permuted. In other words, it is a measure of misfit when all useful information about a parameter is discarded.

**Table 1.**
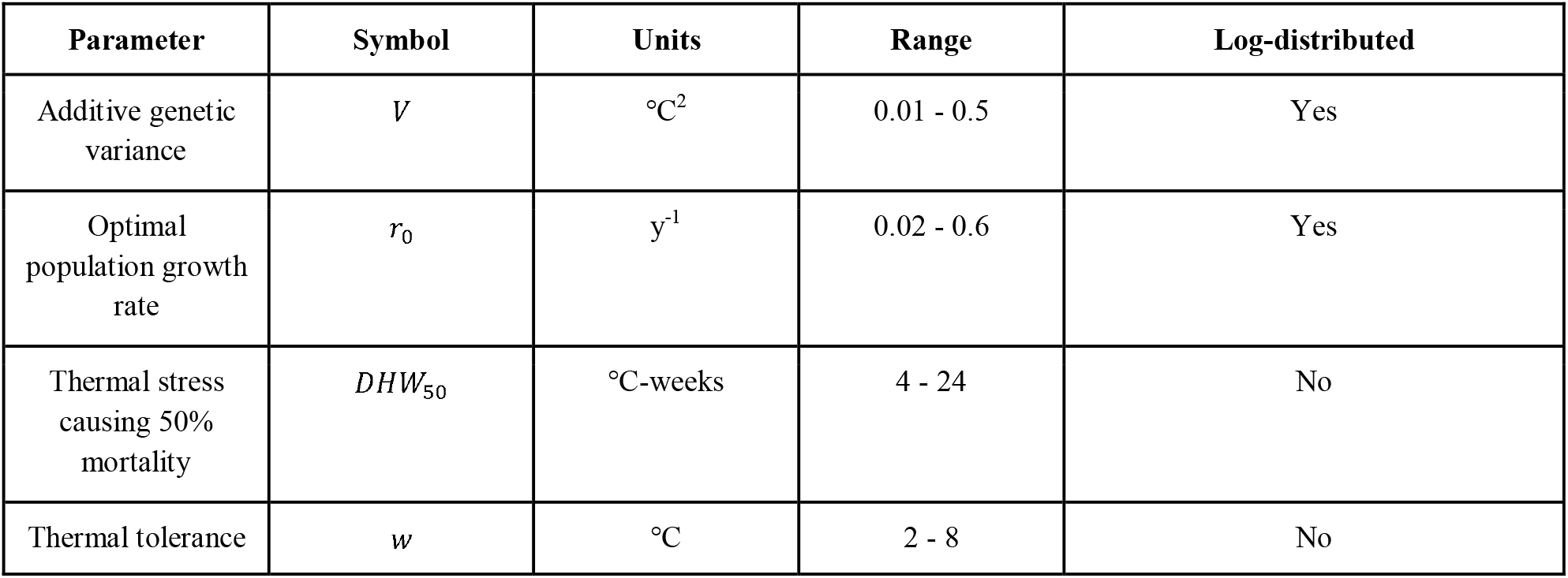
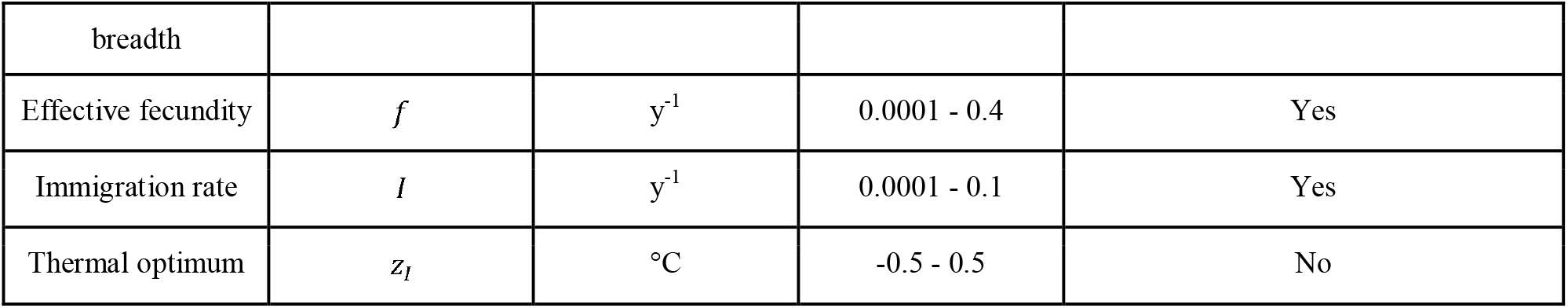
Parameter ranges used in the sensitivity analysis. Parameters marked as ‘Yes’ for ‘Log-distributed’ are distributed logarithmically within these ranges as they vary by more than 1 order of magnitude; all other parameters are distributed linearly.

## Results

Despite differences in temperature variability, inter-site differences in coral cover decline in our model system were very minor, with the standard deviation among sites generally being less than 10% of the mean. This indicates that multidecadal warming trends (which were very similar among all sites) dominated the simulated coral cover decline. Due to these small differences, all further analyses are based on the mean coral cover decline across all sites.

In general, the most important model parameters in determining 21^st^ century coral decline were the optimal population growth rate *r*_*0*_ and the additive genetic variance *V* (figure 1). In this model system, both parameters were first-order controls on coral cover decline under all emissions scenarios (figures S3-4). The coral thermal optimum breadth *w* was consistently the least important parameter. There were some differences in the ranking of these parameters under different emissions scenarios; in particular, the relative importance of additive genetic variance was considerably higher under the most rapid climate change (figure S4). However, in general, these patterns were consistent across all three emissions scenarios tested.

**Figure 1.**
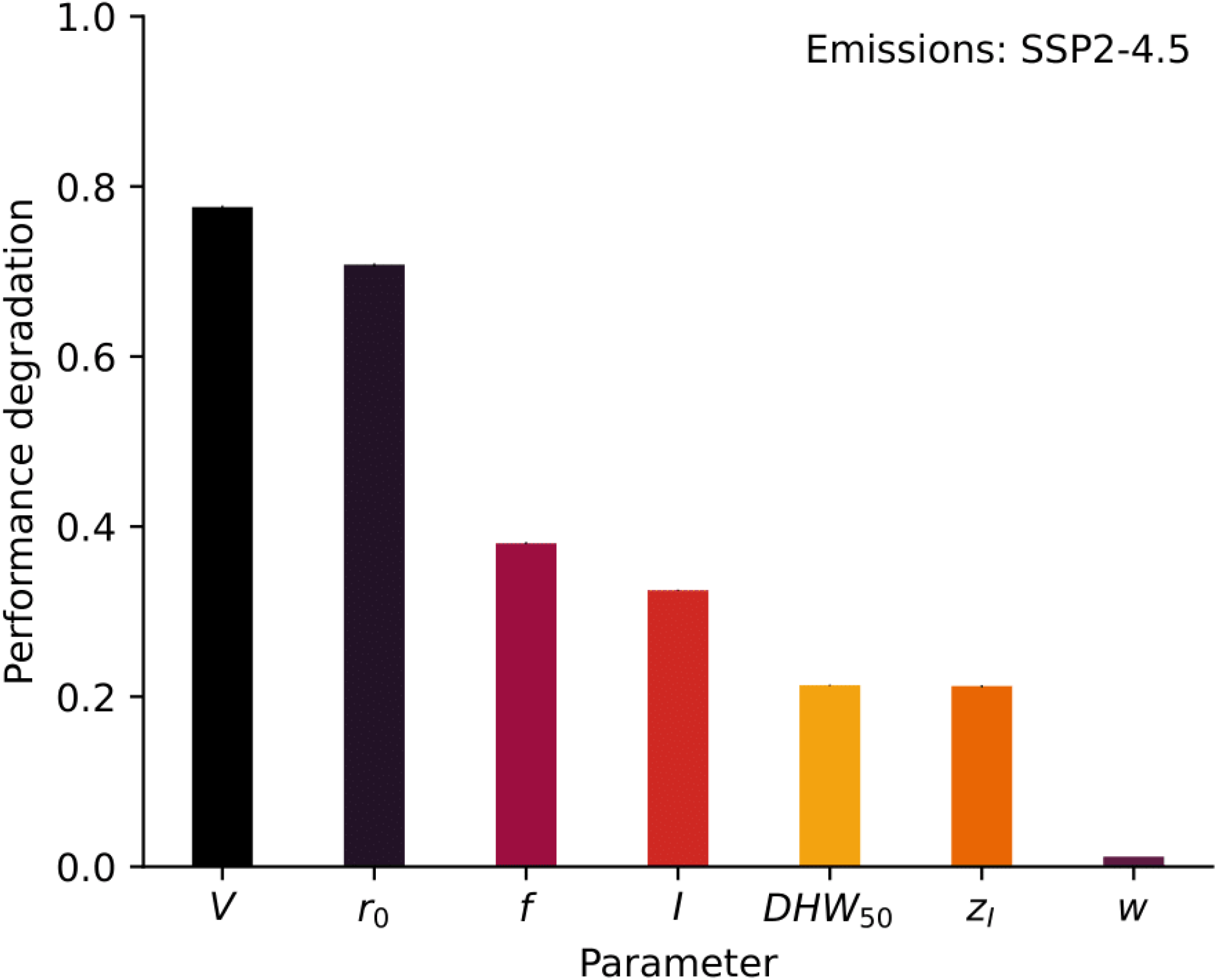
Sensitivity of coral cover decline (the relative change in coral cover from the start of the simulation to the decade with the lowest mean coral cover) to each of the seven model parameters listed in Table 1 based on a random forest regressor and permutation importance, under the SSP2-4.5 emissions scenario. Small vertical lines show the uncertainty in permutation importance based on 10 permutations (the uncertainty is low due to the performance of the random forest regressor in capturing the behaviour of the model system).

The range of biological and ecological parameters described in table 1 result in diverse behaviours under future projected climate change. Without exceptionally high fecundity and/or immigration, both coral growth rate and additive genetic variance must be on the higher end of the plausible range for high coral cover to persist under projected warming (figure 2), specifically growth rate and additive genetic variance exceeding c. 0.1 y^-1^ and c. 0.1 °C^2^ respectively.

**Figure 2.**
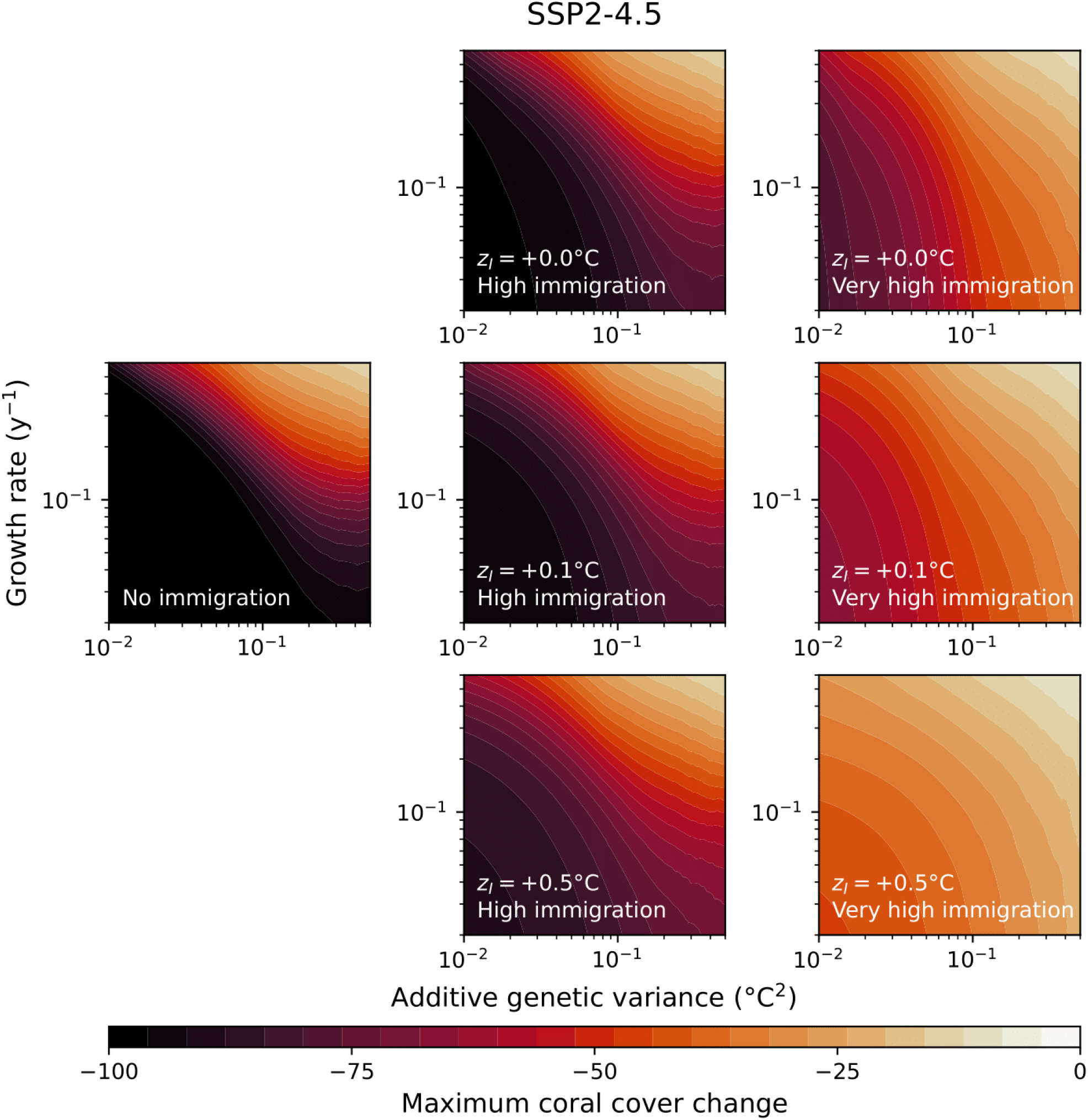
21^st^ century coral cover decline as a function of additive genetic variance (x) and optimal population growth rate (y) under no immigration (left), high immigration (z_l_ = 0.01 y^-1^, centre), and very high immigration (z_l_ = 0.1 y^-1^, right), with immigrant larval thermal optima equal to (top), 0.1 C above (centre) and 0.5 C above (bottom) the population mean thermal optimum. For all cases, DHW_50_ = 8 C-weeks, w = 4 °C, and = 0.01 y^-1^. Bottom-left and top-left panels are missing as immigrant larval thermal optima cannot be modified when there is no immigration.

While immigrant larvae adapted to higher temperatures did have a positive effect on coral persistence, this effect was only apparent at high immigration rates, equivalent to a contribution of around 1% coral cover per year (figure 2). High fecundity and thermal stress tolerance increased the range of *r*_0_ *-V* parameter space consistent with coral persistence (figure 3), but *r*_0 ∼_ 0.1 y^-1^ and *V* ∼ 0.1 °C^2^ broadly remained as threshold values below which persistence was less likely. Higher values of fecundity allowed coral populations to persist under lower growth rates, but had less of an effect on population persistence under low genetic variance.

**Figure 3.**
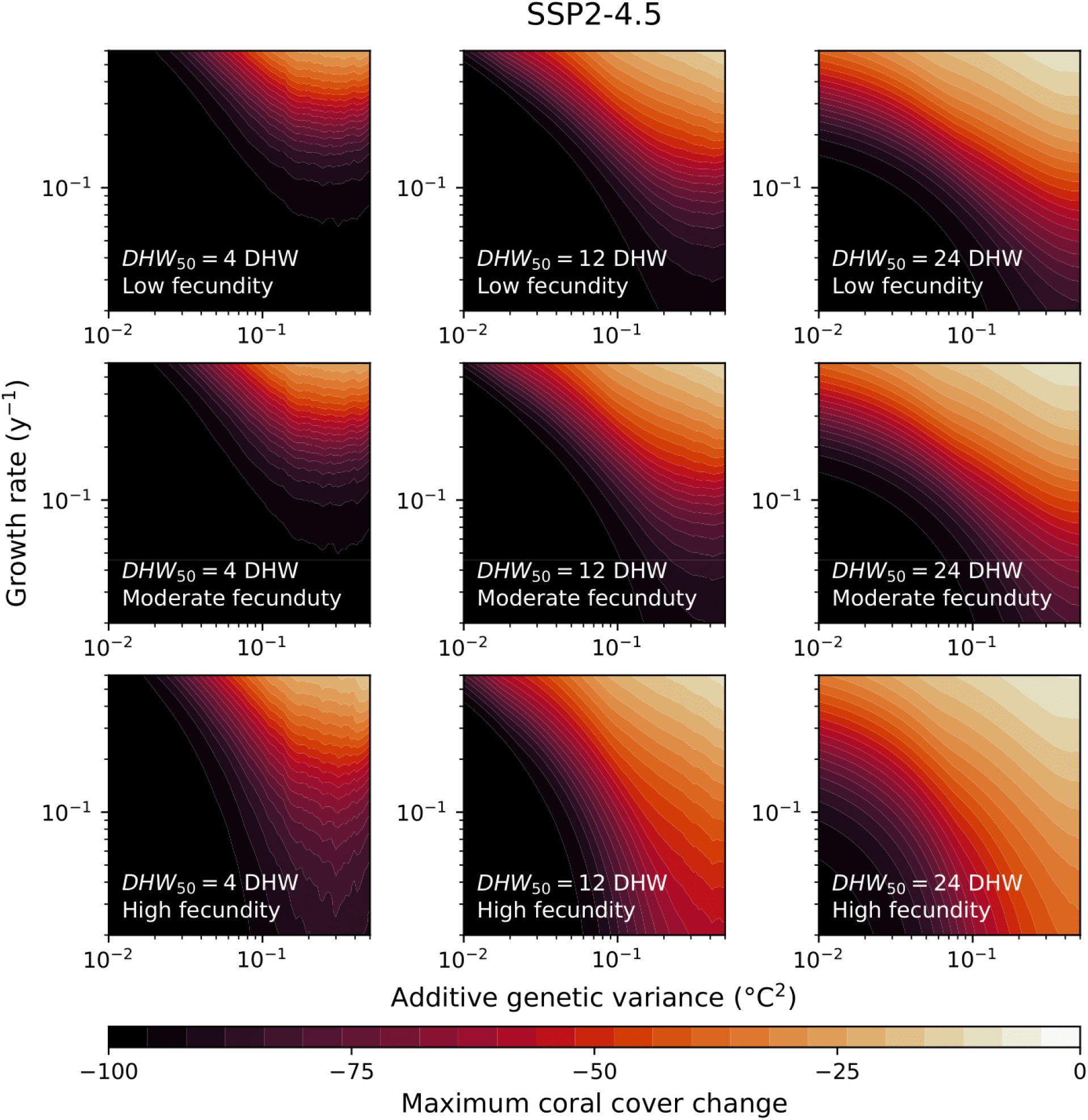
21^st^ century coral cover decline as a function of additive genetic variance (x) and optimal population growth rate (y) under low (4 DHW, left), high (12 DHW, centre), and very high (24 DHW, right) thermal stress tolerance, with low (f = 0.001 y^-1^, top), moderate (f = 0.01 y^-1^, centre) and high (f = 0.1 y^-1^, top) fecundity. For all cases, I = 0.001 y^-1^, z_I_ = 0 °C, and w = 4 °C.

Although the thermal stress tolerance *DHW*_50_ was less important in determining the relative coral cover decline metric (figure 1), it was one of the most important parameters in setting the timing and absolute value of this minimum (figure S5-6), and coral cover variability over sub-decadal timescales. Simulated populations with low thermal stress tolerance reached more extreme coral cover minima more quickly, whereas those with higher tolerance did not reach their minima until the end of the century (figure S7-8). However, in our simulations, the effects of thermal stress tolerance were most pronounced in the early-to-mid 21^st^ century; coral cover at the end of the 21^st^ century had a low sensitivity to thermal stress tolerance, instead again being determined by the optimal population growth rate and additive genetic variance (figure S9).

## Discussion

### Population growth rate and adaptive capacity are critical for long-term persistence

Reefs worldwide are projected to experience more frequent severe bleaching events and increased mortality rates throughout this century (57,58). Identifying the main factors affecting coral persistence under different climate scenarios can help establish priorities for conservation and management efforts.

Here we used an eco-evolutionary model forced by projected future ocean temperatures for Hawai□i, and found that population growth rate and additive genetic variance play a critical role in long-term coral persistence under climate change, with high thermal tolerance alone being insufficient to maintain acceptable levels of coral cover.

There has been considerable interest in determining thermal stress thresholds for corals (11,59,60). However, our simulations suggest, over the course of the entire 21st century, that optimal coral population growth rate (a function of linear extension rate and population size structure) and the additive genetic variance (a function of thermal optimum heritability and phenotypic variability) are critical parameters in setting coral population decline under future climate change. Coral populations with the greatest growth and adaptation potential have the best chance of maintaining high coral cover throughout the century, even if they experience greater mortality in response to marine heatwaves. High additive genetic variance for coral thermal optima (above c. 0.1 °C^2^) allows corals to adapt to higher temperatures over multiple generations through selection, whereas higher growth rates (exceeding around 0.1 y^-1^) provide a mechanism by which coral populations can recover from increasingly frequent mortality events. This is particularly important under more pessimistic warming scenarios, where rapid adaptation is the only mechanism by which coral populations can persist (figure S4).

Genetic diversity in corals is influenced by a combination of factors, including population size (61), sexual reproduction (51), larval dispersal (62), somatic mutations (63), and environmental heterogeneity (64,65). There is natural variability in coral heat tolerance within individual reef environments, which has been observed at values both above and below the threshold value of c. 0.1 °C^2^ identified by this study (43). However, given the existence of well-connected thermal environments within reefs (53) that corals can be locally adapted to (9,64), reefs that are geomorphologically and hydrodynamically diverse (and therefore more likely to harbour thermally distinct sub-reef scale environments) may provide an excellent foundation for coral persistence. Quantifying thermal and genetic variability within coral reefs, and integrating these quantities into reef-scale models (66) may therefore be useful for predicting potential coral ‘refugia’.

### Thermal tolerance is important, but not the full story

While our sensitivity analysis found that population growth rate and adaptation potential are the main drivers of long-term coral population persistence in our model, the thermal stress tolerance (*DHW*_*50*_) was nevertheless important. Since our coral decline metric is based on decadal averages, short-term variability in coral cover is averaged out. However, this variability is considerably higher under lower thermal stress tolerance, increasing the risk that the population could be wiped out entirely by a severe marine heatwave. Indeed, coral taxa with high population growth rates such as most acroporids and pocilloporids have experienced the greatest mortality in response to recent mass bleaching events as they tend to have lower thermal stress thresholds (1,11). Sensitivity analyses considering the *absolute* minimum coral cover (and therefore, potentially, extirpation risk), rather than the relative change in coral cover, as in figure 1, also find a greater importance for *DHW*_50_ (figures S6).

Furthermore, our simulations assume that population size structure remains constant, which is unrealistic (67,68). Particularly if juvenile corals are disproportionately impacted by bleaching mortality, the resulting reduction in the population growth rate (equation 9) would increase the likelihood of a catastrophic population decline. Figure 4 shows simulated coral cover trajectories for two ‘model’ coral species: a branching coral with a high growth rate but low thermal tolerance, and a massive coral with a low growth rate but high thermal tolerance (36). In our base case, the massive coral experiences a slower population decline and a similarly slow recovery. The branching coral reaches lower levels of absolute coral cover in the early-to-mid 21^st^ century, but effectively recovers to higher levels of coral cover than the massive coral in the second half of the century. However, with a 10-fold increase in the median coral colony size, as could perhaps occur with heavy mortality among juvenile corals, coral cover reaches close to zero and never effectively recovers before 2100.

**Figure 4.**
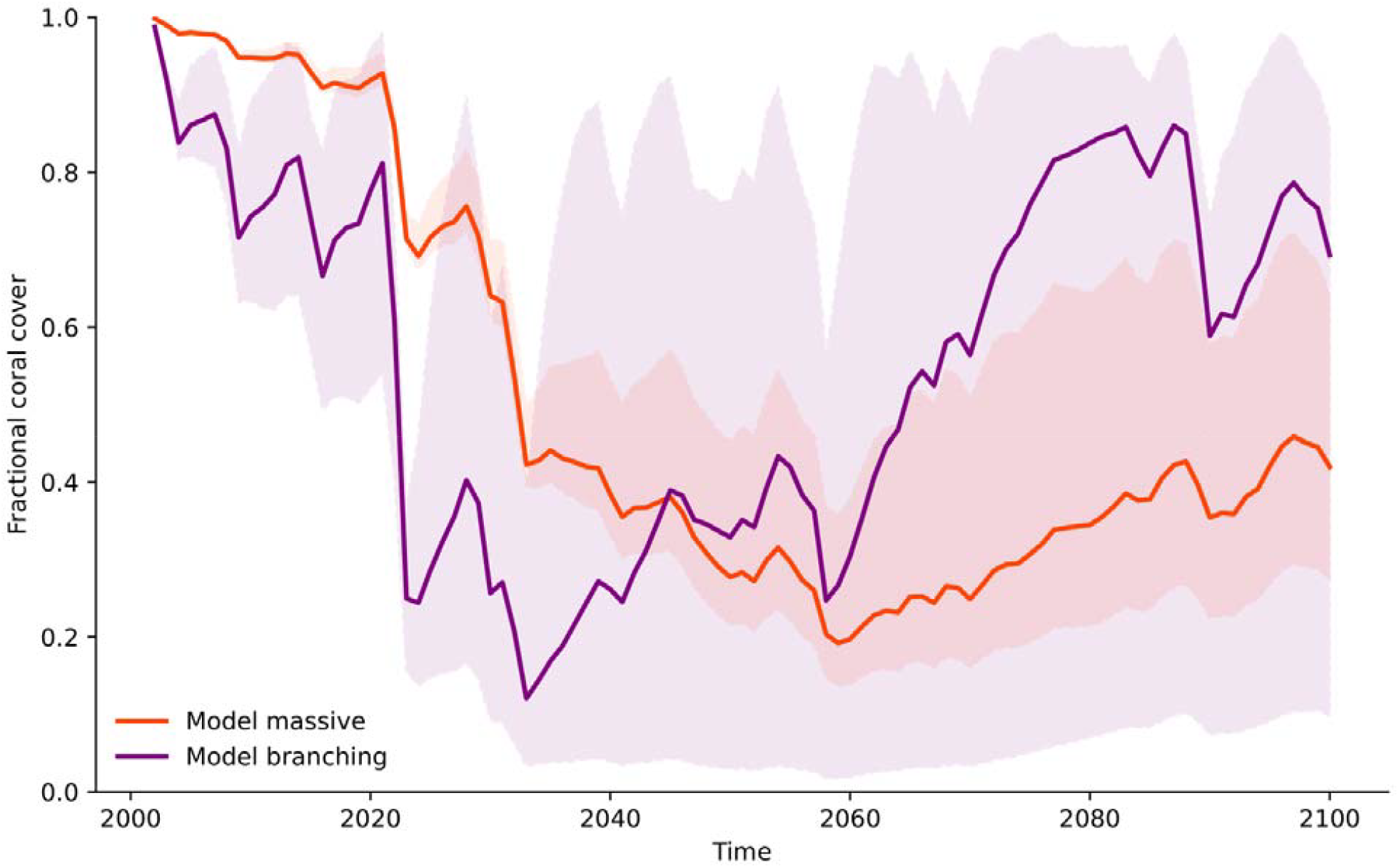
Simulated coral cover trajectories for a model massive (ε =1 cm y^-1^, w=4 °C, f=0.1 y^-1^, DHW_50_ = 15 °C-weeks) and branching (ε =5 cm y^-1^, w=2 °C, f=0.008 y^-1^, DHW_50_ = 5 °C-weeks), based broadly on (36). V=0.1 °C^2^, I=0.001 y^-1^ and z_I_ =0 °C in both cases. Bold line shows the case where the colony size distribution factor is set to the median found in (37), with the shaded region representing the median colony size being increased/decreased by an order of magnitude.

While corals with a low thermal stress tolerance are therefore more vulnerable to experience population collapse, particularly during the mid-to-late 21^st^ century, high coral thermal tolerance does not in itself guarantee that coral populations will persist in response to anthropogenic climate change, instead simply delaying coral cover decline (e.g. figure S7). This is not surprising, since most reefs will regularly experience severe heat stress relative to modern baselines by the end of the century under all reasonable emissions scenarios (58). Indeed, assuming coral populations survive, our simulations suggest that the thermal stress threshold is only of minor importance for setting coral cover at the end of the 21^st^ century (figure S8). In line with previous theoretical studies (10,21), more thermally vulnerable coral taxa could potentially outcompete slower growing, stress tolerant counterparts later this century (figure 4).

As a result, the magnitude of the population response over timescales of <50 years may be dominated by coral thermal tolerance, whereas genetic variance and population growth rate become increasingly dominant post-2050. Therefore, while high thermal tolerance is indeed important in avoiding coral population collapse, it is not sufficient for coral populations to thrive over longer timescales.

### Restoration efforts must focus on building genetic diversity

Previous work has highlighted the potential importance of connectivity, specifically to warmer upstream environments, in supporting coral persistence (21,33). While not the most important parameter in our analyses, immigrant larvae with thermal optima 0.5 °C above the local thermal optima had a positive impact on coral persistence (figure 2), although the required immigration rates were unrealistically high for most long-distance inter-reef connections through natural larval dispersal.

Corals with warmer thermal optima could also be introduced into a reef through transplantation (69) or assisted larval settlement (70). The necessary immigration rates with a thermal optimum offset of +0.5 °C (figure 2) are unlikely to be realistic at scale but, with an offset of +1 °C, (artificial) immigration rates of 0.001 - 0.01 y^-1^ could improve the ability for some corals to persist, boosting the 21^st^ century minimum coral cover by several absolute percentage points (figure S10, see also (71)). This strategy is less effective for populations with low growth rate (due to weaker recovery capacity), and both high genetic variance and growth rates (since these do well even without intervention). Conforming to our results, (71) suggested that restoration approaches focused on building genetic variance may outperform those based solely on introducing heat-tolerant genotypes. However, the required artificial immigration rate of at least 0.001 y^-1^ is nevertheless very high, equivalent to adding around 0.1% of the total reef coral cover per year. Most coral restoration efforts to date have focused primarily on selecting and propagating thermally tolerant corals to enhance short-term persistence (42,72–75). However, we propose that, while thermal tolerance is important, maintaining or enhancing genetic diversity may play an even more critical role in the resilience of coral populations.

While we avoided modelling specific species for generality, our results can be transferred to species and environments of interest, as our analysis suggests that long-term coral cover trajectories largely depend on growth rate and additive genetic variance, as long as thermal tolerance is sufficient to survive the early-to-mid 21^st^ century. While additive genetic variance is largely subpopulation-dependent (64,76,77), growth rate varies significantly between species (78), indicating that the likelihood of persisting under rapid warming may, to some extent, be species-specific. Particularly in subpopulations with greater additive genetic variance, fast-growing species such as many acroporids, montiporids and pocilloporids could plausibly experience a recovery post-2050 (figure 5). For example, *Montipora capitata*, one of the most abundant coral species in Hawai□i, combines fast growth, high thermal resilience, and significant genetic diversity across the Hawaiian archipelago (79) and even at smaller spatial scales, such as Kāne□ohe Bay (9,51,80). Based on our simulations, we can infer that this species has a comparatively positive long-term outlook under future warming, with the major proviso that this relies on those populations weathering bleaching events over the coming decades. Conversely, slow-growing massive corals such as those in the *Porites* genus generally exhibit high thermal tolerance (81,82), which allows them to resist initial bleaching events and maintain structural integrity when more sensitive species decline. Although *Porites* species often display moderate to high genetic diversity in some regions (83,84), this varies geographically and may not fully compensate for their slower demographic recovery rates. As such, their recovery from anthropogenic climate change may be slower and less effective in the longer run.

**Figure 5.**
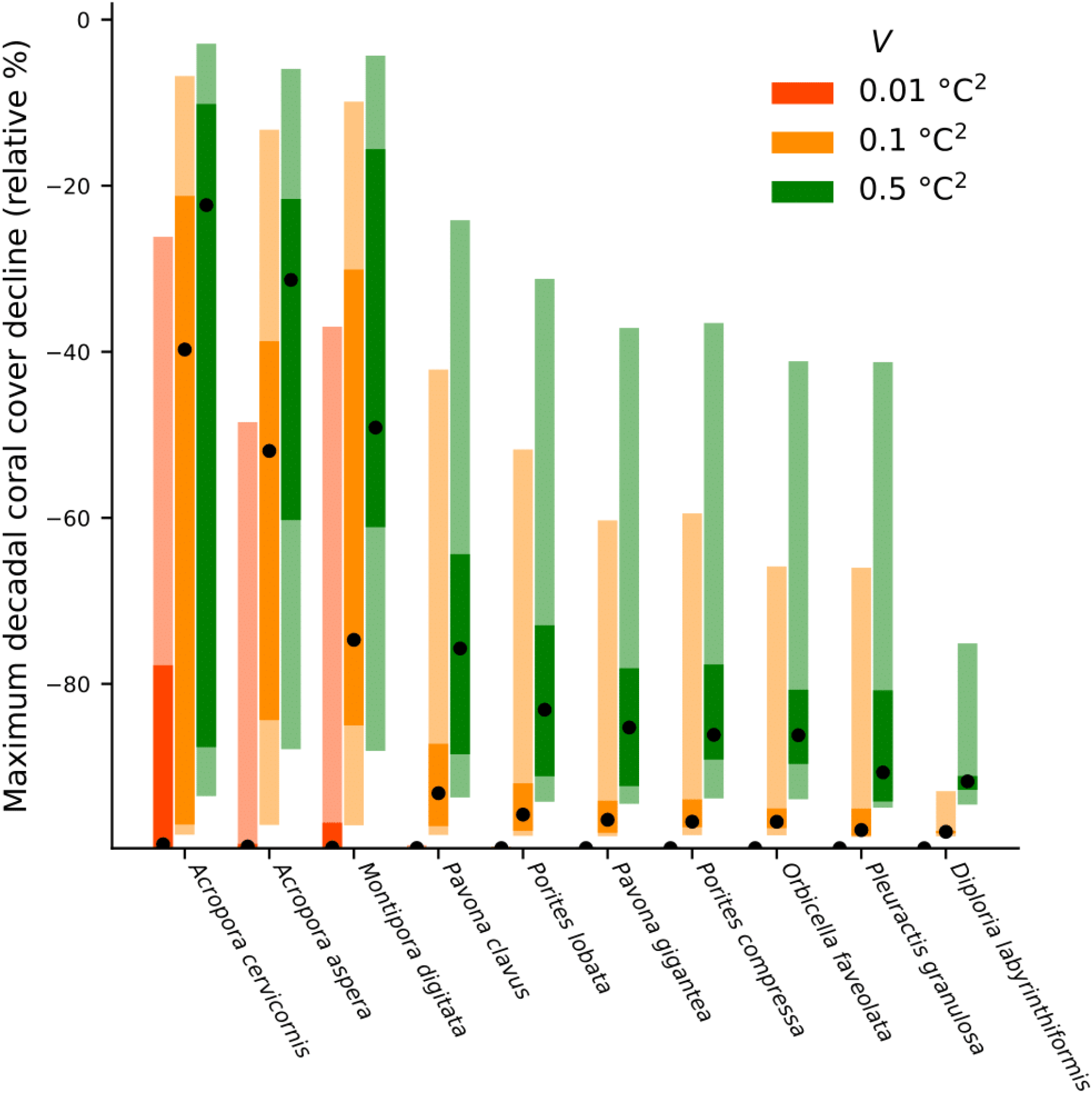
Maximum decline in coral cover (decadal averages) over the 21^st^ century based on the 5-95^th^ percentile (bars, pale extensions representing a factor of 10 increase/decrease in the median coral size as in figure 4) and median (dots) linear extension rates for various coral species from the Coral Traits Database (78), under low (red), moderate (orange) and high (green) additive genetic variance (V). Simulations assume DHW_50_ = 8 °C-weeks, w = 4 °C, = 0.01 y^-1^, I = 0.001 y^-1^, and z_I_ = 0 °C, with size structure set to the average across populations investigated by (37). We emphasise that non-growth parameters are not tuned to named species here - we name species solely to emphasise potential consequences of inter- and intraspecific variation in growth rate.

The model framework used in this study was necessarily simple to explore such a large parameter space. We model the additive genetic variance as being constant; while genetic diversity has been maintained through past environmental disturbances (52), this assumption may be violated for sites with low standing genetic diversity, poor connectivity, and/or low effective population size. Our model also assumes that coral population size structure remains fixed, when it is in reality dynamic (68), affecting the effective population growth rate. While we simplistically incorporated uncertainty in size structure in analyses (figures 4-5), future studies could use size-structured models to more comprehensively model resulting population dynamics. We do not model interspecies interactions, but these interactions could affect coral population dynamics depending on the effects of climate change on competing species (32). The parameterisation of connectivity in this study is simplistic, assuming that larval immigration remains static, whereas it would in reality likely decline in tandem with upstream coral population sizes. It is also possible that mechanisms not included in our model may play a role in supporting population persistence. For example, factors such as coral acclimatization (85) and the interactions between corals and their dinoflagellate symbionts (Symbiodiniaceae) play a crucial role in the holobiont -encompassing both the animal and its microbial community - response to ocean warming (86,87). These symbiotic relationships can be particularly important in coral responses to temperature and susceptibility to bleaching (88). This study uses Hawaiian reefs as a case study. Hawai’i provides a uniquely valuable natural laboratory, as its corals already experience thermal and pH conditions that are not expected to occur on most reefs until later in the century (89). As such, Hawaiian reefs offer a valuable lens through which to explore persistence potential under future climate regimes, but may underestimate the thermal heterogeneity found at other locations, particularly at higher latitudes, where greater temperature variability suggests that thermal tolerance breadth may play an especially critical role in determining persistence.. Finally, temperature data used (34) are subject to model and scenario uncertainty, and are provided at a resolution that cannot resolve reef-scale temperature variability,

## Conclusion

Even under optimistic emissions scenarios, anthropogenic climate change will inevitably outpace the rate of adaptation for practically all corals. As a result, within our simple eco-evolutionary model system, coral population persistence was largely determined not just by coral thermal tolerance, but by adaptive capacity and population growth rate. While the coral thermal tolerance appears to be particularly important in terms of setting the early response of coral populations to warming and the threat of population collapse in the short term, population growth rate and genetic variance become increasingly important for population persistence later in the 21^st^ century. This conclusion was robust under both optimistic and pessimistic emissions projections.

Fast-growing coral taxa often have lower thermal tolerance, and have been disproportionately affected by early bleaching events (11). However, these groups also have the greatest capacity to recover from mass mortality events, and our simulations suggest that this advantage may increase post-2050, and vice versa for slower growing, stress-tolerant species (figure 4). We therefore caution against the assumption that coral species that have shown greater resilience in the early 21^st^ century will fare similarly in coming decades.

Adaptive capacity, largely determined by genetic diversity, is a critical factor in coral persistence that should guide management through coral restoration efforts. As coral reefs continue to decline, global interest in restoration initiatives has grown significantly (100, but see 101). However, a major challenge is that restored populations often exhibit lower genetic diversity than natural counterparts, primarily due to a limited number of donor colonies or minimal genetic variation among them (92,93). This reduction in diversity increases the risk of genetic swamping, and our results demonstrate that the establishment of low-diversity coral subpopulations through coral restoration is not a sustainable strategy, even if they appear to experience lower mortality in the short term. To date, however, most coral restoration efforts have focused primarily on thermal tolerance as the key trait to enhance persistence (72,73,94–96), whereas long-term reef resilience requires maintaining not only genetic but also functional diversity across species, ensuring that a range of ecological niches are sustained (97). The dual importance of growth rate and coral thermal tolerance in setting coral population resilience, combined with a broad trade-off between growth rate and thermal tolerance at a species level (1,11,36) further underlines the importance of diversity in reef restoration approaches, if restored reefs are to survive the challenges of the coming century.

## Data and code availability

All data and code required to reproduce the results and figures in this manuscript have been made available to reviewers as electronic supplementary materials. Upon acceptance, these data will be made available in a Zenodo repository, referenced in this section.

## Acknowledgements

N.S.V.-V. is supported by the NOAA Climate and Global Change Postdoctoral Fellowship Program, administered by UCAR’s Cooperative Programs for the Advancement of Earth System Science (CPAESS) under award #NA21OAR4310383 and #NA23OAR4310383B. M.R.S. is supported by National Science Foundation NSF BioOCE#2048457. L.C.M. acknowledges support by the National Science Foundation under awards 2233983 and 2443233. L.C.M and R.J.T. acknowledge support by the National Oceanic and Atmospheric Administration NOS-ONMS-MOA-2022-008 (Amendment 001)/12868. K.F. acknowledges support of the Uehiro Center for the Advancement of Oceanography. The technical support and advanced computing resources from University of Hawaii Information Technology Services—Cyberinfrastructure, funded in part by the National Science Foundation CC* awards #2201428 and #2232862, are gratefully acknowledged. Finally, we thank the Marine Ecological Theory Lab at the Hawai□i Institute of Marine Biology for their thoughts and comments on this manuscript.

