## Supplementary Materials for "Growth rate and adaptive capacity, not just thermal tolerance, are critical for long-term coral persistence under climate change"

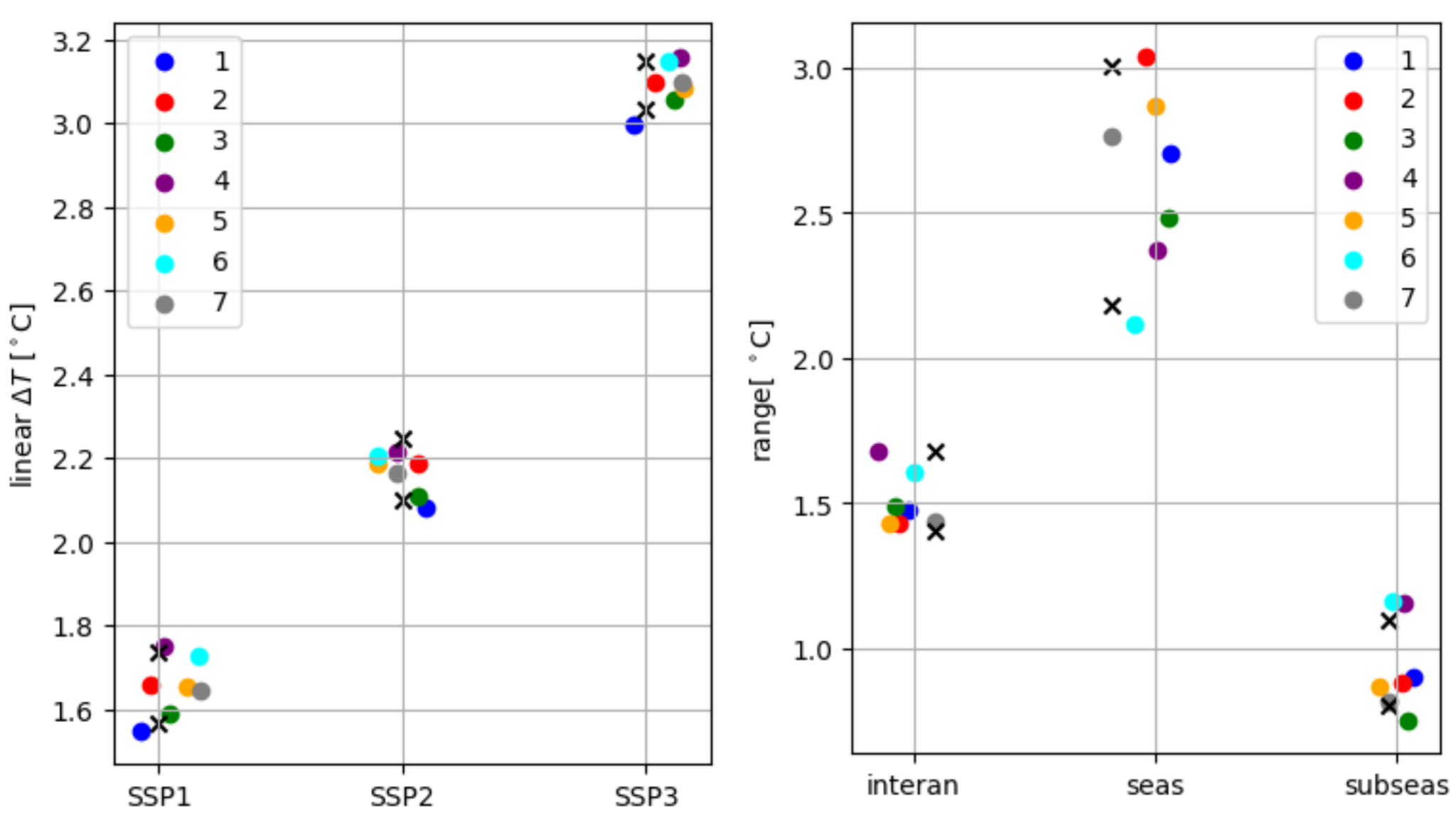

*Figure S1:* (Left) *Linear simulated warming trend over the 21st century under SSP1-2.6, SSP2-4.5, and SSP3-7.0 at each site (see figure S2 for map reference).* (Right) *Magnitude of interannual (>380 days), seasonal (300 - 380 days), and subseasonal (30 - 300 days) variability at each site. Crosses show the 5-95 percentiles across all coastal points within the model domain.*

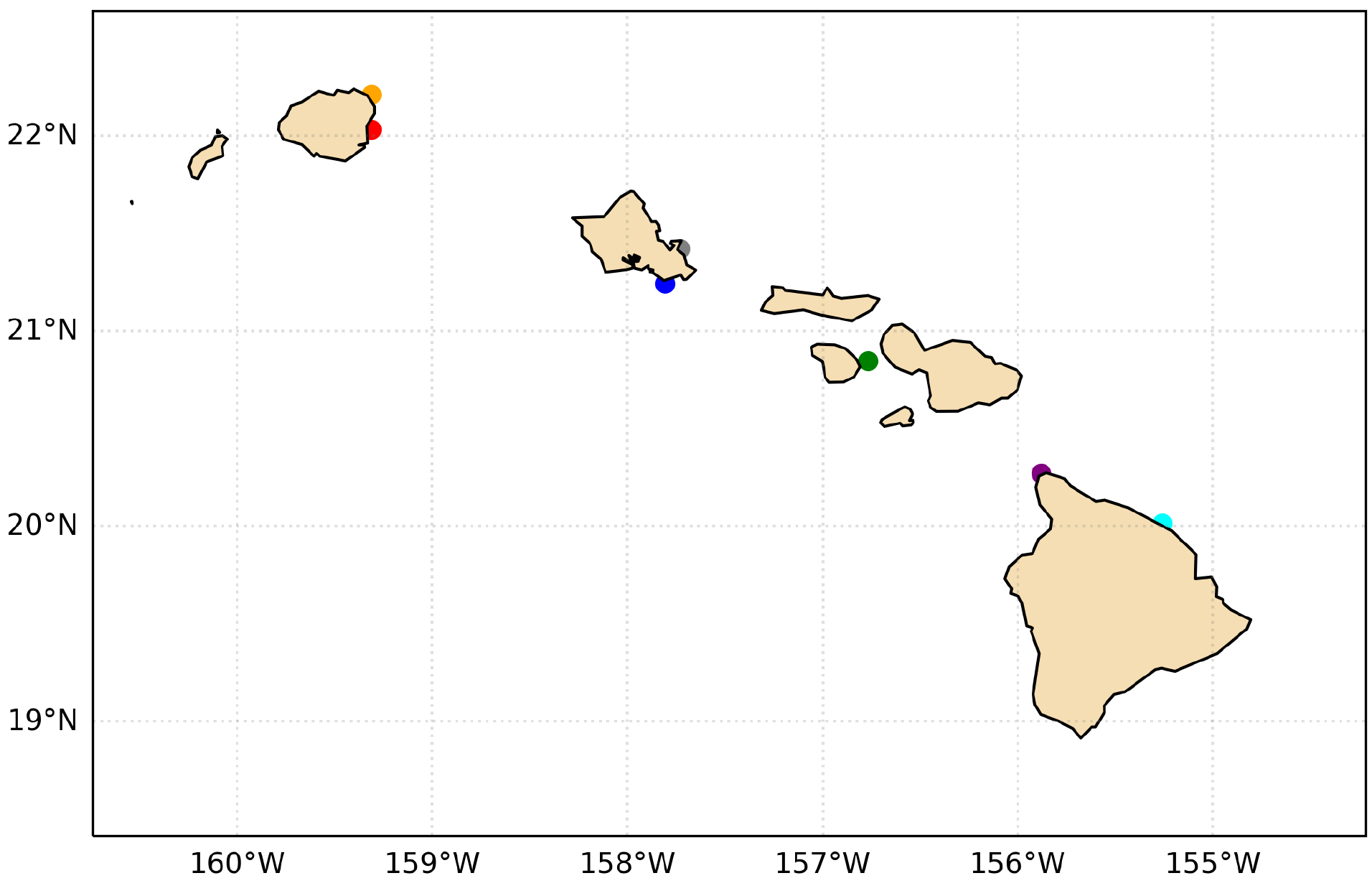

*Figure S2: Map of the Main Hawaiian Islands showing the locations of the seven chosen modelled locations .*

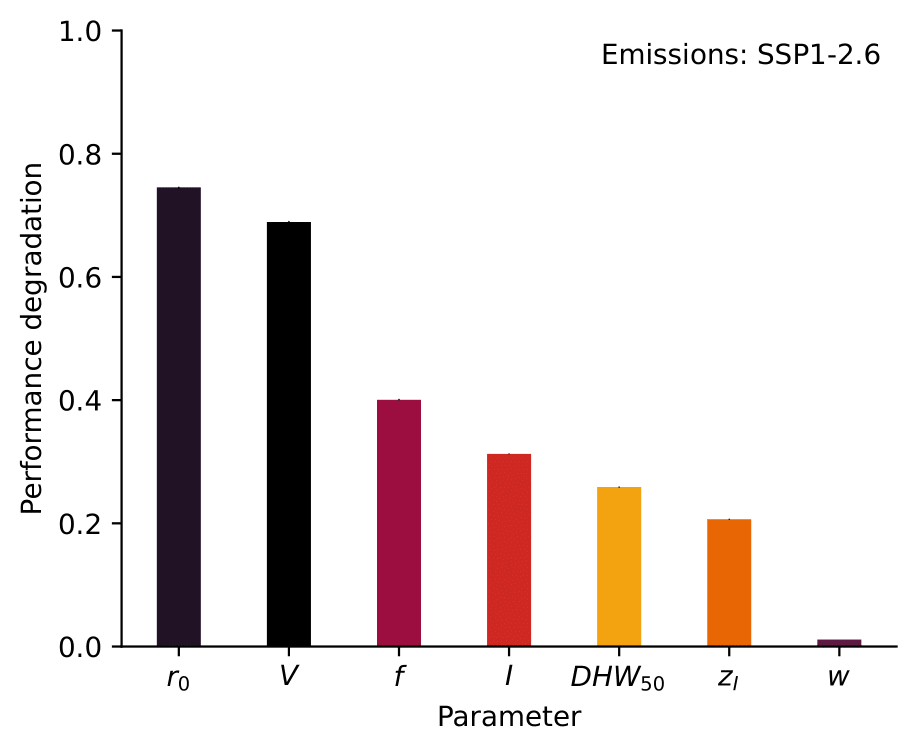

*Figure S3: Sensitivity of* ***coral cover decline*** *to each of the seven model parameters listed in Table 1 based on a random forest regressor and permutation importance, as in figure 1, but for the SSP1-2.6 emissions scenario. Vertical lines show the uncertainty in permutation importance based on 10 permutations (note that this is difficult to see as the uncertainty is extremely low due to the well-behaved model system).*

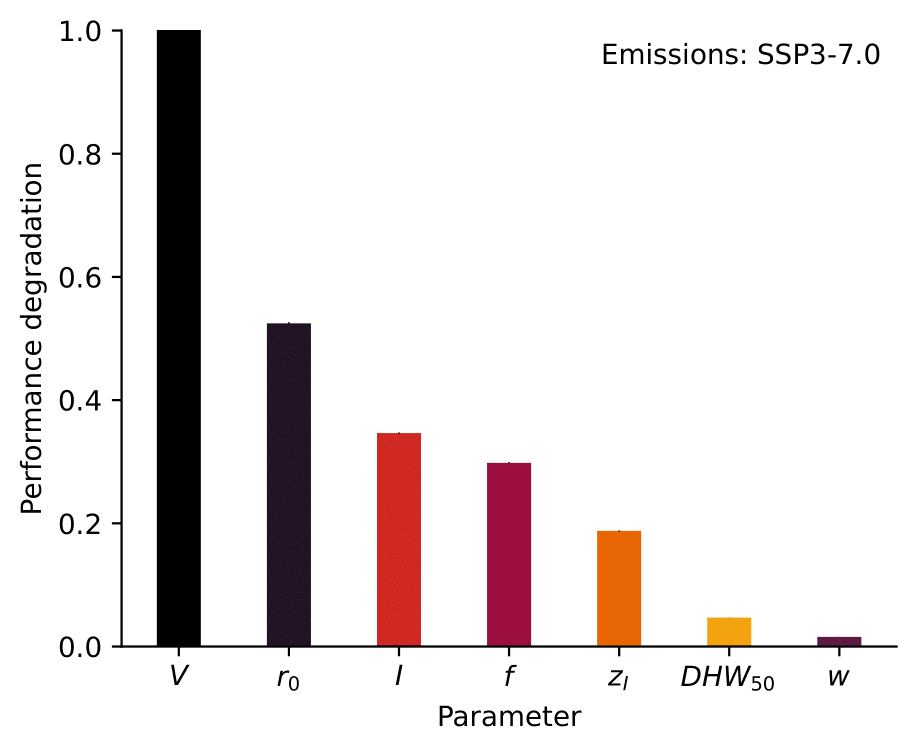

*Figure S4: Sensitivity of* ***coral cover decline*** *to each of the seven model parameters listed in Table 1 based on a random forest regressor and permutation importance, as in figure 1, but for the SSP3-7.0 emissions scenario. Vertical lines show the uncertainty in permutation importance based on 10 permutations.*

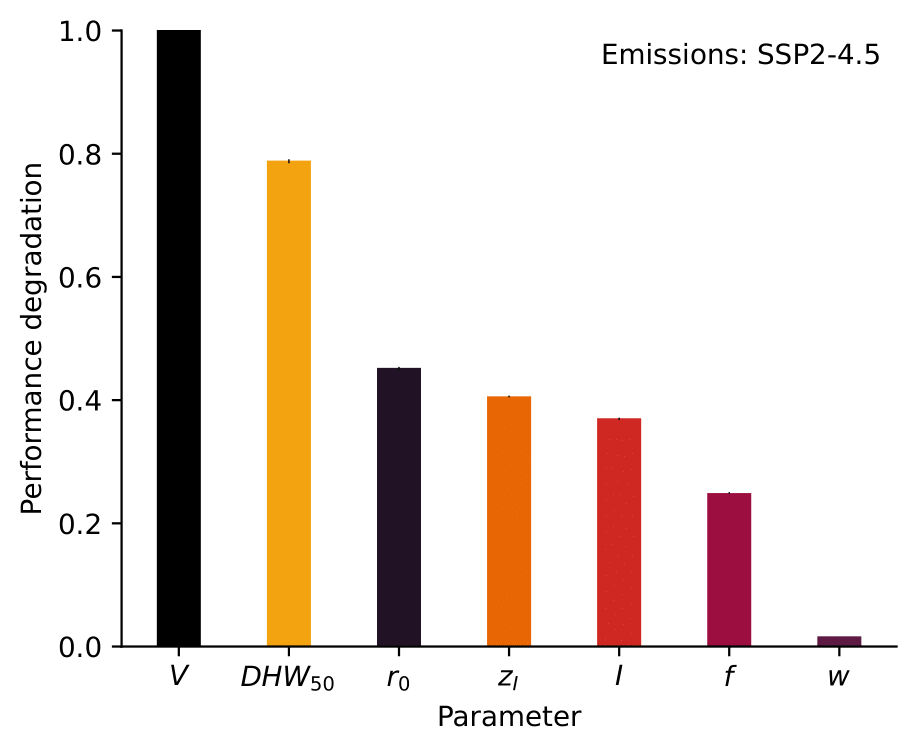

*Figure S5: Sensitivity of the* ***decade during which minimum coral cover is reached*** *to each of the seven model parameters listed in Table 1 based on a random forest regressor and permutation importance, under the SSP2-4.5 emissions scenario. Vertical lines show the uncertainty in permutation importance based on 10 permutations.*

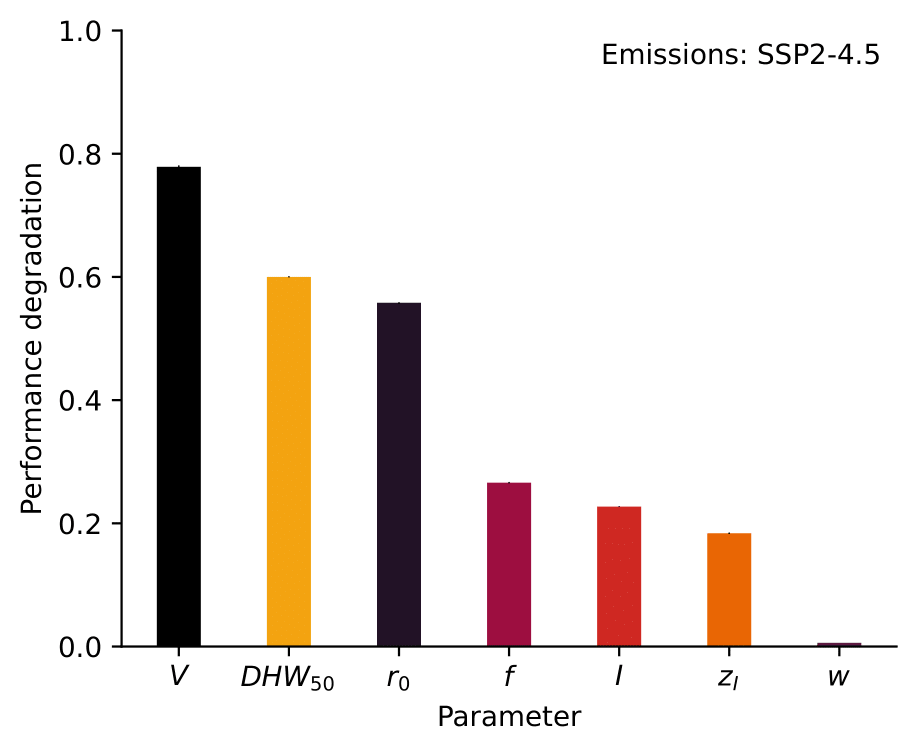

*Figure S6: Sensitivity of the* ***absolute minimum coral cover attained throughout the 21st century (without decadal averaging)*** *to each of the seven model parameters listed in Table 1 based on a random forest regressor and permutation importance, under the SSP2-4.5 emissions scenario. Vertical lines show the uncertainty in permutation importance based on 10 permutations.*

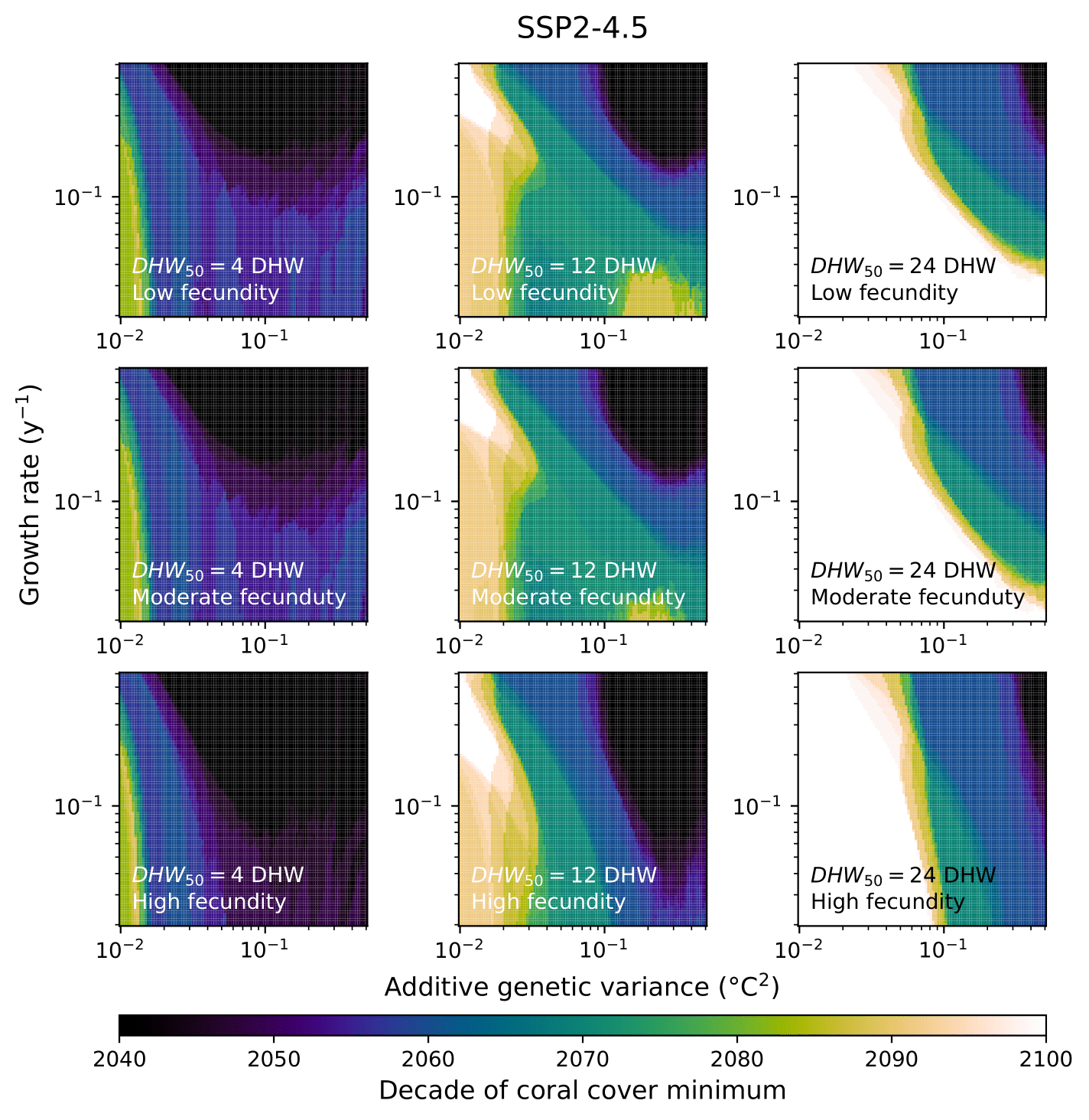

*Figure S7: Decade during coral cover switches from decline to recovery, as a function of additive genetic variance (x) and optimal population growth rate (y) under low (4 DHW, left), high (12 DHW, centre), and very high (24 DHW, right) thermal stress tolerance, with low (*$f=0.001$ *y^-1^, top), moderate (*$f=0.01$ *y^-1^, centre) and high (*$f=0.1$ *y^-1^, top) fecundity. For all cases,* $I=0.001$ *y^-1^,* $z_{I}=0$ °*C, and* $w=4$ °*C.*

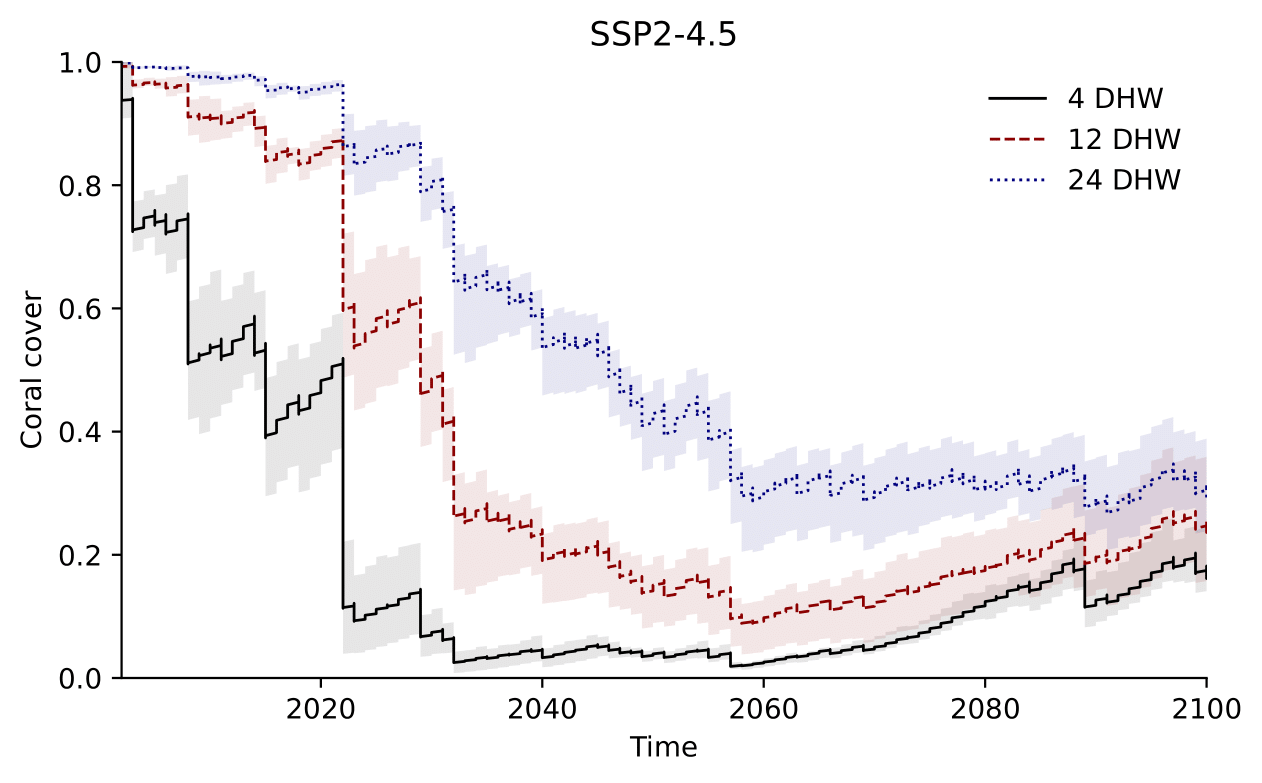

*Figure S8: Three example time-series of coral cover under low (4 DHW, black solid), high (12 DHW), red dashed), and very high (24 DHW, blue dotted) thermal tolerance. For all cases,* $V=0.1$ *K^2^,* $r_{0}=0.1$ *y^-1^,* $f=0.01$ *y^-1^,* $I=0.001$ *y^-1^,* $z_{I}=0$ *C, and* $w=4$ *C. Shaded regions span the minimum and maximum coral cover across the 7 sites.*

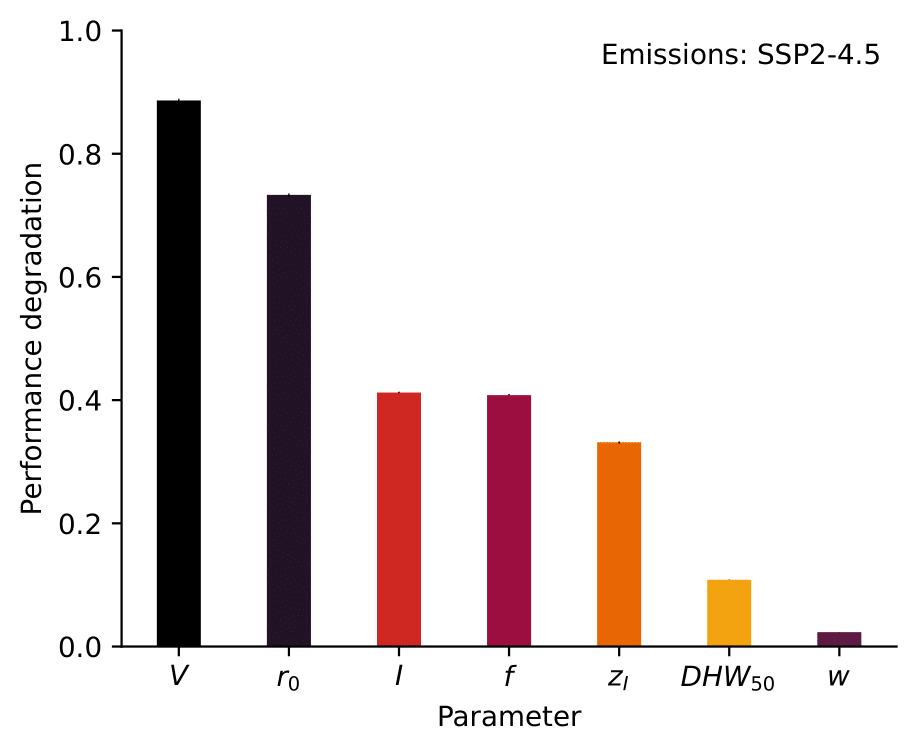

*Figure S9: Sensitivity of the* ***relative change in coral cover from the start of the simulation to 2090-2100*** *to each of the seven model parameters listed in Table 1 based on a random forest regressor and permutation importance, under the SSP2-4.5 emissions scenario. Vertical lines show the uncertainty in permutation importance based on 10 permutations.*

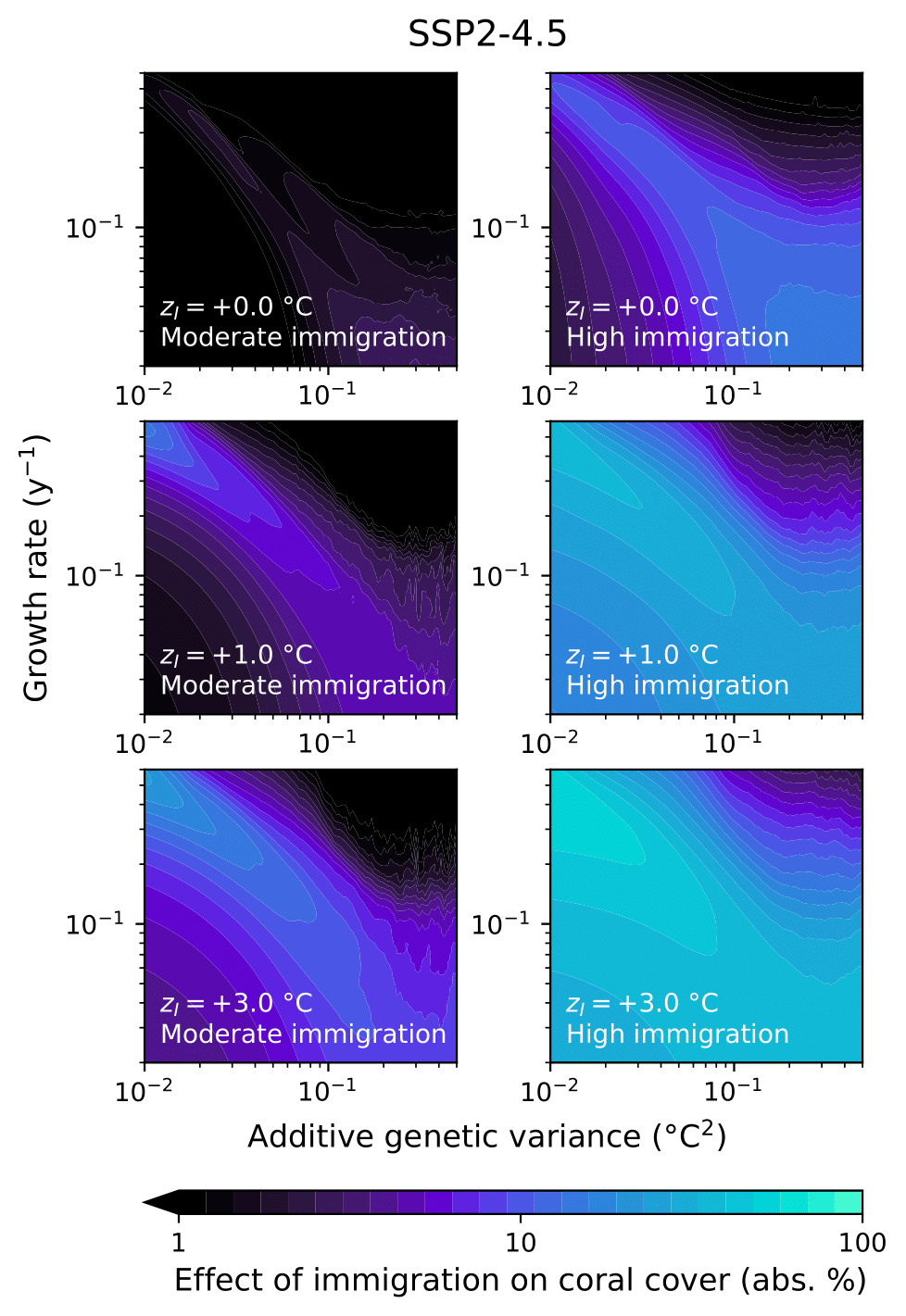

*Figure S10: Effect of larval immigration, quantified as the absolute difference in minimum coral cover versus zero immigration, as a function of additive genetic variance (x) and optimal population growth rate (y) under moderate immigration (*$z_{I}=0.001$ *y^-1^, centre), and high immigration (*$z_{I}=0.01$ *y^-1^, right), with immigrant larval thermal optima equal to 0* °*C (top), 1* °*C above (centre) and 3* °*C above (bottom) the population mean thermal optimum. For all cases,* $DHW_{50}=8$ °*C-weeks,* $w=4$ °*C, and* $f=0.01$ *y^-1^.*
